# Lipid nanoparticle composition for adjuvant formulation modulates disease after influenza virus infection in QIV vaccinated mice

**DOI:** 10.1101/2024.01.14.575599

**Authors:** Sonia Jangra, Alexander Lamoot, Gagandeep Singh, Gabriel Laghlali, Yong Chen, Tingting Yz, Adolfo García-Sastre, Bruno G. De Geest, Michael Schotsaert

## Abstract

Adjuvants can enhance vaccine effectiveness of currently licensed influenza vaccines. We tested influenza vaccination in a mouse model with two adjuvants: Sendai virus derived defective interfering (SDI) RNA, a RIG-I agonist, and an amphiphilic imidazoquinoline (IMDQ-PEG-Chol), TLR7/8 adjuvant. The negatively charged SDI RNA was formulated into lipid nanoparticles (LNPs) facilitating the direct delivery of a RIG-I agonist to the cytosol. We have previously tested SDI and IMDQ-PEG-Chol as standalone and combination adjuvants for influenza and SARS-CoV-2 vaccines. Here we tested two different ionizable lipids, K-Ac7-Dsa and S-Ac7-Dog, for LNP formulations. The adjuvanticity of IMDQ-PEG-Chol with and without empty or SDI-loaded LNPs was validated in a licensed vaccine setting (quadrivalent influenza vaccine or QIV) against H1N1 influenza virus, showing robust induction of antibody titres and T cell responses. Depending on the adjuvant combination and LNP lipid composition (K-Ac7-Dsa or S-Ac7-Dog lipids), humoral and cellular vaccine responses could be tailored towards type 1 or type 2 host responses with specific cytokine profiles that correlated with protection during viral infection. The extent of protection conferred by different vaccine/LNP/adjuvant combinations was examined against challenge with the vaccine-matching strain of H1N1 influenza A virus. Groups that received either LNP formulated with SDI, IMDQ-PEG-Chol or both showed very low levels of viral replication in their lungs at five days post virus infection. LNP ionizable lipid composition as well as loading (empty versus SDI) also skewed host responses to infection, as reflected in the cytokine and chemokine levels in lungs of vaccinated animals upon infection. These studies show the potential of LNPs as adjuvant delivery vehicles for licensed vaccines and illustrate the importance of LNP composition for subsequent host responses to infection, an important point of consideration for vaccine safety.

## Introduction

After decades of research into influenza virus vaccines, the respiratory virus is still a major global health concern causing thousands of cases of severe medical illness in humans every year. Several licensed influenza vaccine candidates, including recombinant, inactivated and split influenza vaccines, have been developed and eventually licensed for use in the human population^1,2^. Despite the availability of licensed vaccines, the need to update and vaccinate people every year remains a challenge as the circulating influenza viruses can escape host immunity provided by antibodies that target the immunodominant but ever-changing antigenic sites on the hemagglutinin (HA) protein^3–5^. Vaccination against both seasonal Influenza A (IAV) and influenza B (IBV) has been effective in controlling virus-related disease severities. However, the protection provided by humoral immunity induced by these vaccines is reported as antigenically constricted and short term. Moreover, the vaccine-induced neutralizing antibody titres drop over time, rendering the immunity less effective against an antigenically different strain of virus in the subsequent seasons^6–8^. Therefore, to combat the need of a seasonal vaccine, a better cost-effective approach is required in vaccine development which can provide a broader and long-term immune response that lasts for multiple seasons.

Inactivated split virus vaccines, including trivalent inactivated vaccines (TIV) and quadrivalent inactivated vaccines (QIV), are most commonly used influenza vaccines. TIV comprises of two IAV strains, one each of H1N1 and H3N2 subtype, and one IBV component, while QIV consists of two IAV and two IBV strains components (representing the Yamagata and Victoria linages)^9,10^. These vaccines can induce strain-specific antibody responses with high serum IgG levels *in vivo* but are poor inducers of cell-mediated immunity and therefore, provide limited protection against antigenically drifted virus strains. Due to the continuous acquisition of mutations in antigenic sites of the viral hemagglutinin, the protective effect of currently licensed seasonal influenza virus vaccines is time confined.

Novel vaccine concepts that aim at inducing broader, long-lasting immunity against influenza virus infection are based on enhancing vaccine- induced B and T cell responses which can recognize multiple antigens from vaccine components, with special focus on targeting the conserved viral epitopes. While natural infection typically results in the induction of type 1 responses, characterised by Th1 and in BALB/c mice class switching to serum IgG2a antibodies to clear viral infection^11–13^, inactivated split virus influenza vaccines typically induce high IgG1 levels correlating with Th2-type immune response^14–16^. Therefore, many studies, including our recently published study^17^, have been focusing on combining commercially available vaccines with specific adjuvants to specifically direct responses to IgG2a or IgG1, hence inducing Th1/Th2 responses^18–21^. Eventually, an efficiently balanced humoral response with enhanced T cell activation post-vaccination is desired to be protective.

Lipid nanoparticles (LNPs) are non-viral vectors that are widely used in formulating vaccines and/or adjuvants to enhance their antigenicity and improve immune responses^22,23^. LNPs have already shown promising outcomes in formulating antigen-encoding mRNA, such as SARA-CoV-2 mRNA vaccines^24^. These mRNA vaccine-LNP formulations have also successfully demonstrated the role of LNP-based vaccine platform for an efficient induction of humoral and cell-mediated immunity. Moreover, LNPs can also be used for formulating molecular adjuvants, such as RIG-I or TLR agonists, and facilitate uptake by actively phagocytosing innate immune cell subsets^25^. Nevertheless, the composition of LNP is crucial to achieve the optimal uptake by innate immune cells and efficient humoral responses. A typical LNP consists of four main components: an ionizable lipid, a phospholipid, cholesterol moiety and polyethylene-glycol (PEG)-lipid. The ionizable lipids consist of ionizable positively charged lipids that can effectively interact with negatively charged mRNA molecules. Phospholipids and cholesterol provide structural stability to LNPs and facilitate endosomal escape, thus enhancing efficient delivery of mRNA into the cytosol of cells. The PEG lipids prolong the circulation of LNPs consisting of vaccines/adjuvants in circulation by increasing their half-life. Additionally, the surface molecules of LNPs can also be modified to target specific innate immune cells and facilitate uptake for efficient antigen presentation^23,25–27^. Overall, LNPs present as an efficient *in vivo* vaccine-adjuvant delivery system.

In this study, we investigated and compared the efficiency of two LNP formulations, consisting of different ionizable cationic lipids, in inducing both humoral and cell mediated immune responses in a mouse model receiving a single shot of QIV from 2018-19 influenza season, with and without adjuvants, individually or in combination. Specifically, we used an *in vitro* transcribed Sendai virus defective-interfering RNA^17,28,29^ (SDI-RNA; a RIG-I agonist; negatively charged and hence encapsulated into LNPs) and an amphiphilic imidazoquinoline conjugate^17,30^ (IMDQ-PEG-Chol; a modified TLR7/8 agonist with enhanced safety profile and lymph node-draining properties), previously characterized and tested by our groups, as adjuvants. Vaccine-induced antibody and T cell responses were characterized and further correlated with lung cytokine profiles and extent of protection against lethal challenge of a matching strain to H1N1 component of QIV: A/Singapore/GP1908/2015 (IVR-180). We observed adjuvant-specific differences in B- and T- cell responses, which were not only driven by the presence of different adjuvants (SDI RNA and/or IMDQ-PEG-CHOL) but also depended on the type of ionizable/cationic lipid composition of the LNPs.

## Results

### Preparation and characterization of LNPs

Two LNP formulations were prepared by mixing an aqueous solution containing the *in vitro* transcribed SDI-RNA (or SDI) with an ethanolic solution containing (1) ionizable lipids, either K- Ac7-Dsa (comprising a ketal bond) or S-Ac7-Dog (comprising a disulfide bond; chemical structure outlined in supplementary Fig 1), to interact with negatively charged SDI and mediate endosomal escape; (2) cholesterol, for structural stability; (3) dioleoylphosphatidylethanolamine (DOPE) phospholipid, as helper lipids to aid in nanoparticle formation; (4) 1,2-distearoyl-rac-glycero-3- methylpolyethylene glycol (DSG-PEG; 2kDa PEG) to provide steric hinderance and thus avoiding aggregation and promoting mobility *in vivo*. The structure and composition of LNP incorporating SDI-RNA is schematically represented in Fig 1A and B, respectively. The molar ratio of ionizable lipid (K-Ac7-Dsa or S-Ac7-Dog): cholesterol: DOPE: DSG-PEG was chosen to be 50:38.5:10:1.5, based on literature. Empty LNP, that did not contain SDI, were prepared as control and hence referred to as LNP(-). Both LNPs encapsulating SDI (S-Ac7-Dog(SDI) and K-Ac7-Dsa (SDI)) and corresponding control empty LNPs (S-Ac7-Dog(-) and K-Ac7-Dsa (-))were fabricated and characterized for their size and zeta potential (Fig 1C and 1D). While K-Ac7-Dsa(-) and K-Ac7- Dsa(SDI) LNPs showed some differences in their size (78 and 189 nm respectively), the S-Ac7- Dog(-) and S-Ac7-Dog(SDI) LNPs showed similar size distributions (91 and 101 nm respectively) with low polydispersity indices (PDI< 0.25; as shown in Fig 1C) and a positive zeta potential (ZP) around 4-6.5 mV at physiological pH (Fig 1D).

**Figure 1.**
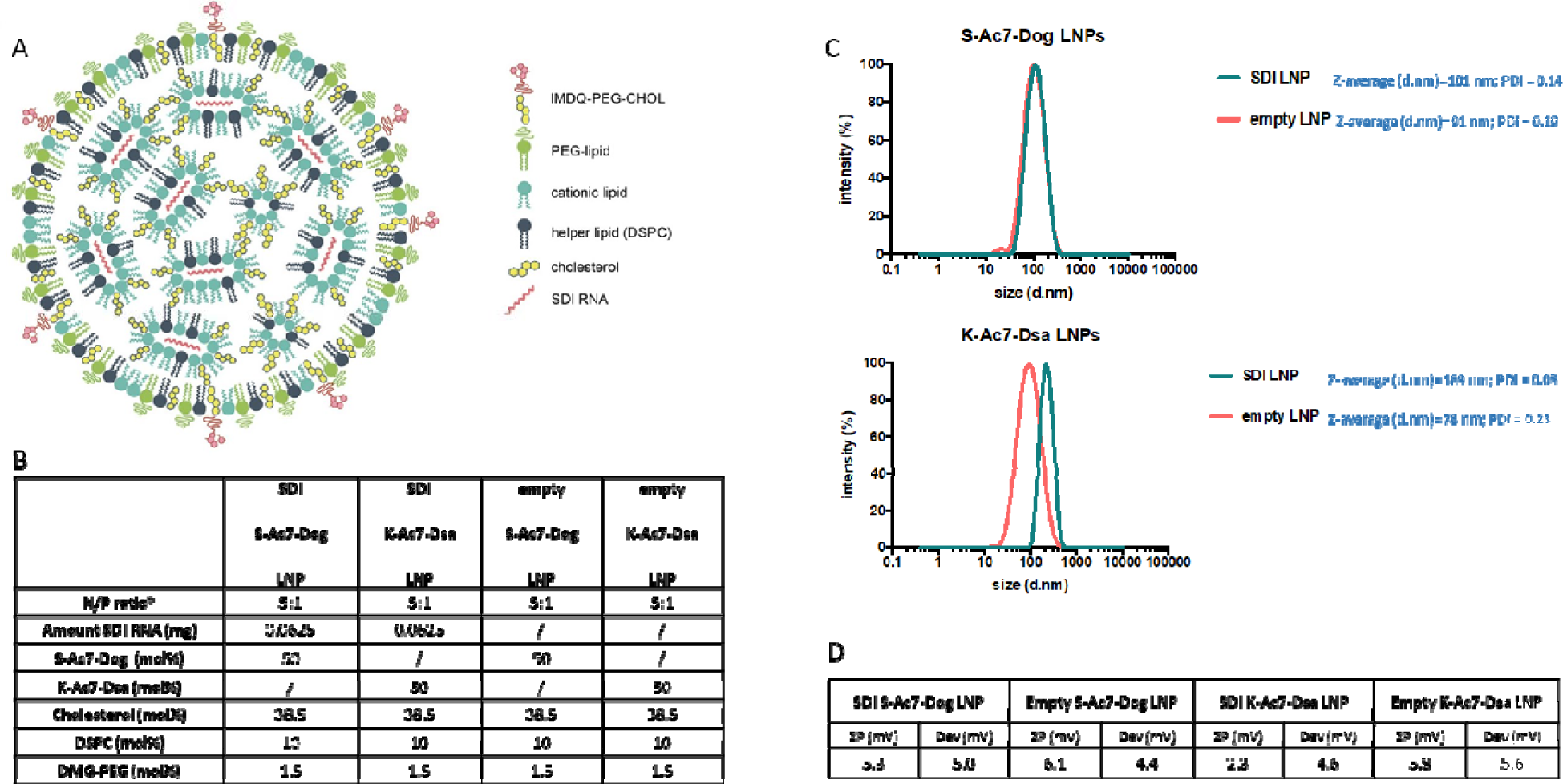
Structure and characterization of LNPs consisting of K-Ac7-Dsa or S-Ac7-Dog lipids. **(A)** Diagrammatic representation of LNP structure encapsulating SDI **(B)** Composition ratio of different components of LNPs, with and without SDI **(C)** Intensity-based size distribution curves measured by Dynamic Light Scattering (DLS) of empty and SDI-incorporating S-Ac7-Dog and K- Ac7-Dsa LNP formulations **(D)** Summarizing table of the LNPs zeta potential measured by Electrophoretic Light Scattering (ELS).

### LNP formulations with adjuvanted QIV define IgG subtype profile, with S-Ac7-Dog LNPs inducing higher antibody titers than K-Ac7-Dsa LNPs

We evaluated the potential of empty and SDI-incorporating S-Ac7-Dog and K-Ac7-Dsa LNPs to adjuvant a licensed quadrivalent influenza vaccine (QIV), with and without combination with IMDQ-PEG-Chol adjuvant, using our previously well-established preclinical vaccination-infection animal model. The study is outlined in Fig 2A. We vaccinated 6-8-week female BALB/c mice with unadjuvanted QIV or in combination with SDI or IMDQ-PEG-Chol or both, while mock animals received PBS. We further tested the adjuvant effect for QIV upon co-administration of SDI and/or IMDQ-PEG-Chol combined with one of the two LNPs containing different cationic lipids (S-Ac7- Dog or K-Ac7-Dsa) as described in previous section. LNPs were either empty (-) or had SDI incorporated. The rationale of this set up is that LNP-formulated SDI can, besides endosomes, also be delivered to the cytosol, thereby promoting more efficient RIG-I mediated innate immune sensing. On the other hand, IMDQ-PEG-Chol is expected to incorporate efficiently in LNPs via its cholesterol moiety. The animals were vaccinated only once via the intramuscular (IM) route. The serum collected 3 weeks post-vaccination was examined for the presence of H1 HA-specific IgG antibodies using enzyme-linked immunosorbent assay (ELISA) for total IgG, IgG1 and IgG2a (Fig. 2b).

**Figure 2:**
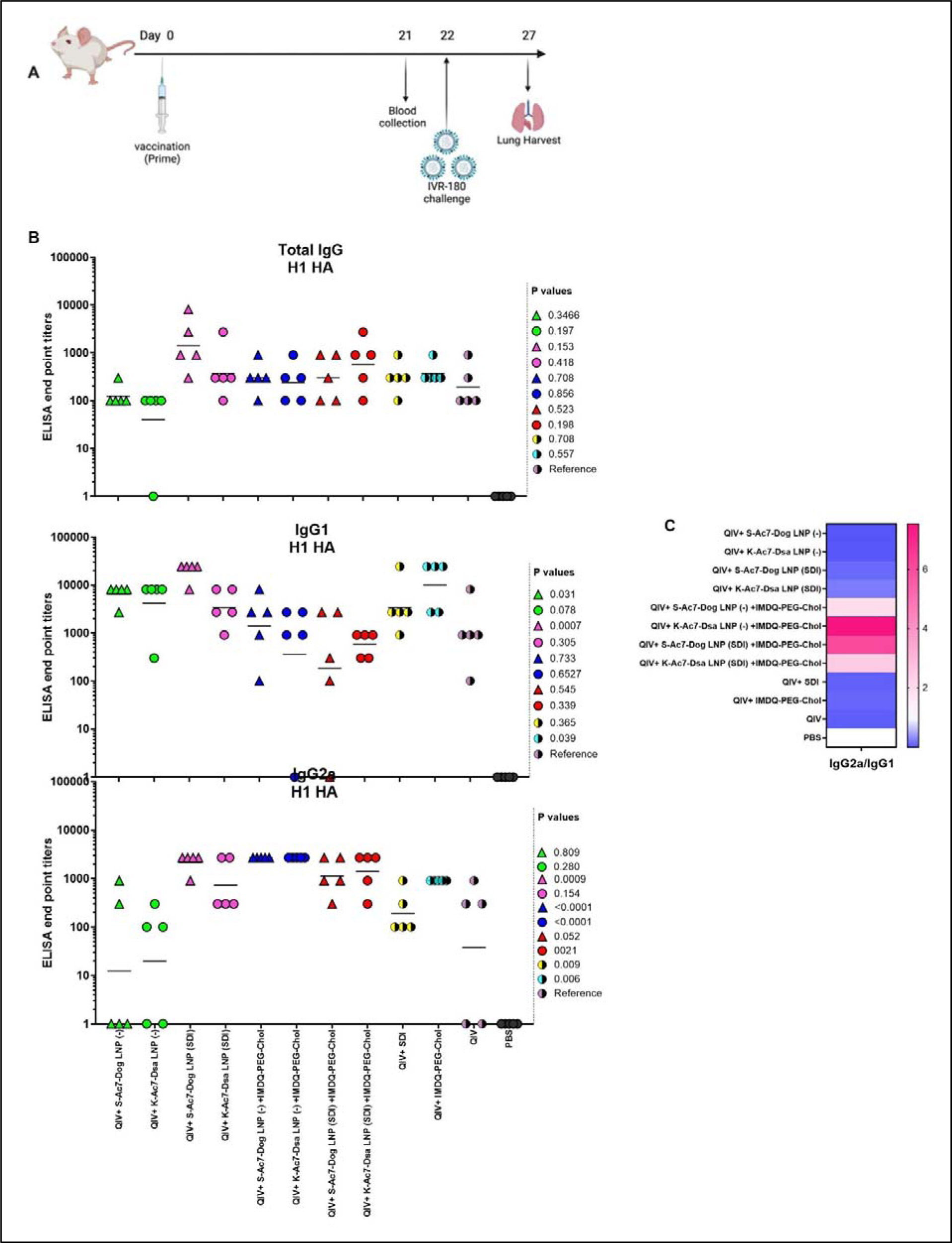
S-Ac7-Dog (- or SDI) and K-ac7-Dsa (- or SDI) in combination with IMDQ-PEG- Chol define IgG subtype profile: (A) Study outline **(B)** Graphs showing ELISA end point titers calculated based on OD450 ELISA values against serum dilutions for total IgG, IgG1 and IgG2a (n=5 per group), represented as geometric mean. The statistical analysis was performed using unpaired T test and the p values shown are calculated in reference to the unadjuvanted QIV group which received neither adjuvant nor LNP formulations. (C) Heatmap showing the ratio of end point titers of IgG2a to IgG1 and represented as geometric mean of IgG2a/IgG1 ratios for all animal in each group.

No antibody titers were detected in the serum from mock PBS group. The group which received unadjuvanted QIV was used as the reference to compare the IgG responses of other groups. QIV formulated with S-Ac7-Dog LNP (-) or K-Ac7-Dsa LNP(-), corresponding to empty S-Ac7-Dog and empty K-Ac7-Dsa LNPs respectively, showed higher IgG1 titers but lowest IgG2a titers compared with the corresponding S-Ac7-Dog LNP(SDI) or K-Ac7-Dsa LNP(SDI) groups (Fig 2B and supplementary Fig 2), thereby illustrating the intrinsic adjuvant effect of LNPs. Additionally, QIV formulated with SDI-incorporated LNPs (S-Ac7-Dog LNP(SDI) and K-Ac7-Dsa LNP(SDI)) can induce a balanced IgG1 and IgG2a antibody response. Overall, the total IgG levels were found similar among all adjuvanted and LNP-formulated groups post-prime vaccination. However, the S- Ac7-Dog LNP(SDI) seemed to be induce slightly higher total IgG slightly compared to K-Ac7-Dsa LNP (SDI) in corresponding SDI ± IMDQ-PEG-Chol combination adjuvanted groups. A similar observation was made for IgG1 antibody titers with S-Ac7-Dog LNPs inducing higher IgG1 titers than respective K-Ac7-Dsa LNP groups. Consistent with our previous findings^17^, IMDQ-PEG-Chol administration, with either of empty or SDI-incorporated S-Ac7-Dog or K-Ac7-Dsa LNPs, skews the antibody responses towards IgG2a with a significant reduction in IgG1 titers (Fig 2C). QIV+SDI and QIV+IMDQ-PEG-Chol groups, with no LNP formulations, were used as control vaccination groups for the study and to correlate with our previous study.

### QIV adjuvanted with IMDQ-PEG-Chol exhibit a better control over virus neutralization when formulated in S-Kc7-Dog LNPs than K-Ac7-Dsa LNPs

Neutralizing antibodies are important defense mechanisms during viral infection. These antibodies bind to the viral antigens and thereby block the viral attachment to cells. As a surrogate for virus neutralizing antibody levels, we performed hemagglutinin inhibition (HAI) assays with post-vaccination sera collected from all vaccinated animals at 3 weeks post vaccination. As shown in Fig 3, the mice that received unadjuvanted QIV showed low HAI titers which were not significantly different from the groups that received QIV formulated in either empty or SDI-containing S-Ac7- Dog or K-Ac7-Dsa LNPs. The group administered with QIV+IMDQ-PEG-Chol, without LNP formulations, showed significantly higher HAI titers compared with unadjuvanted QIV group. HAI titers were significantly higher for the groups that received empty or SDI-containing S-Ac7-Dog in the LNP formulation when combined with IMDQ-PEG-Chol adjuvant. However, the corresponding K-Ac7-Dsa LNP-formulated groups did not show any significant differences in HAI titers compared with unadjuvanted or unformulated QIV group.

**Figure 3:**
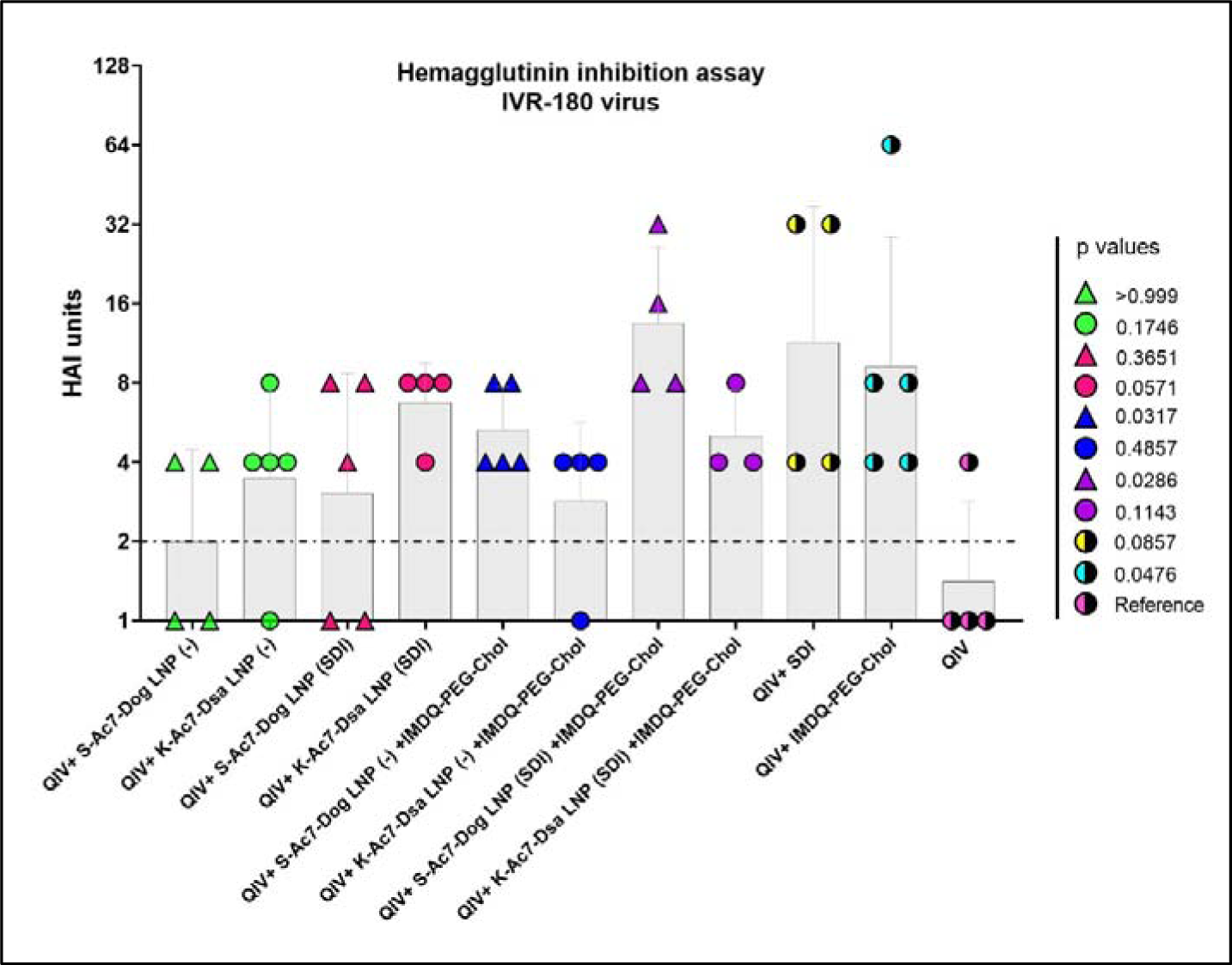
QIV adjuvanted with IMDQ-PEG-Chol exhibits a better control over virus infection when formulated in S-Kc7-Dog LNPs: The sera collected from all vaccinated animals 3 weeks post-vaccination was tested for HAI titers, using 4 HA units of IVR-180 virus. The HAI titers are represented as geometric mean ± geometric SD for n=4 animals per group. The samples with undetectable HAI titers were set as 1 and the limit of detection was set to 2, corresponding to the lowest detectable HAI units. The statistical analysis was performed using two-sided Mann Whitney U test. The p values shown are calculated in reference to the QIV group which received neither adjuvant nor LNP formulations.

### QIV formulated in either of SDI-containing S-Ac7-Dog and K-Ac7-Dsa LNPs efficiently induce T cells responses when combined with IMDQ-PEG-Chol

Helper CD4+ and cytotoxic CD8+ T cells play an important part in vaccination-mediated humoral and cellular responses by facilitating Ig class switch during B cell maturation and direct killing of infected cells, respectively. T cells can recognize foreign antigens presented on the major histocompatibility complex (MHC) molecules on infected cells followed by release of various cytokines, including IFN-γ and IL-4, two major cytokines corresponding to helper type-1 (Th1) and type-2 (Th2) T cells, respectively. IFN-γ and IL-4 can modulate class switching of B cells to IgG2a or IgG1^19^. To study the correlation between the two cytokines and antibody responses in our vaccination model, we examined the release of IFN-γ and IL-4 from splenocytes obtained from the mice 10 days post-vaccination (outlined in Fig 4A), in presence and absence of specific antigen (IVR-180 virus or H1 HA peptide) using enzyme-linked immunosorbent spot (ELISpot) assays.

**Figure 4:**
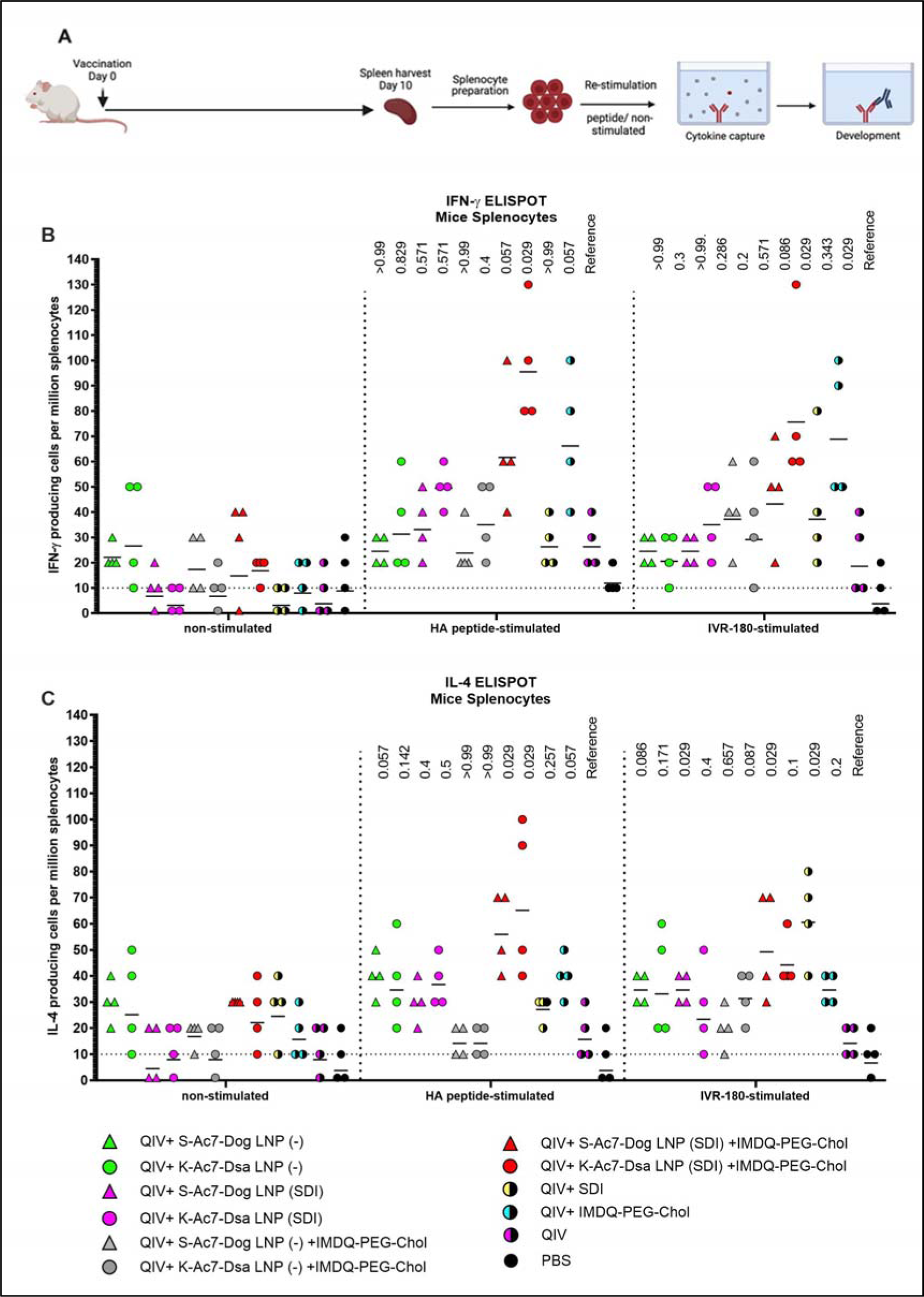
QIV formulated in S-Ac7-Dog(SDI) and K-Ac7-Dsa(SDI) LNPs efficiently induce T cell responses when combined with IMDQ-PEG-Chol: (A) Experiment outline **(B,C)** 6-8 weeks BALB/c mice were vaccinated with QIV with and without IMDQ-PEG-Chol and formulated into S-Ac7-Dog(- or SDI) or K-Ac7-Dsa(- or SDI) LNPs. The spleens were harvested 10 days post- vaccination to examine the T cell activation by IFN-γ **(B)** and IL-4 **(C)** ELIspots, upon restimulation with H1-HA short-overlapping peptides or live IVR-180 (A/Singapore/GP1908/2015 H1N1) virus. The results are represented as IFN-γ or IL-4 producing cells per million splenocytes (geometric mean ± geometric SD) for n=4 animals per group. The cut-off was set to 10 which indicates one spot in any well. The wells with no spots were given the value 1. The statistical analysis was performed using two-sided Mann Whitney U test and the p values shown are calculated in reference to the respective unadjuvanted QIV group.

As shown in Fig 4B and C, antigen-specific IFN-γ (Fig 4B) or IL-4 (Fig 4C) release from the splenocytes was very low in the absence of a stimulant, except for QIV+S-Ac7-Dog(-) or QIV+K- Ac7-Dsa(-) groups suggesting a basal level of non-specific stimulation after vaccination by the empty LNPs. Upon stimulation with specific H1-HA peptide or IVR-180 virus, the number of splenocytes producing antigen-specific IFN-γ as well as IL-4 were found to be higher in the mice that received QIV+IMDQ-PEG-Chol formulated in S-Ac7-Dog(SDI) or K-Ac7- Dsa(SDI) LNPs, suggesting an induction in both Th1 and Th2 immune response. The group administered with QIV+IMDQ-PEG-Chol, without LNP formulations, showed high IFN-γ release compared with unadjuvanted QIV group, consistent with our previous study. Mice that received QIV only were used as a reference for statistical comparison.

### S-Ac7-Dog and K-Ac7-Dsa LNP formulations with SDI and/or IMDQ-PEG-Chol potentiates QIV-mediated protection against a lethal viral challenge with homologous influenza virus

To further corelate B and T cell responses with the extent of protection, all vaccinated and un- vaccinated mice were challenged with 100 times 50% lethal dose of IVR-180 virus (18000 PFU per animal). A single dose of vaccination was found effective in conferring protection from severe morbidity in challenged animals compared with mock-challenged mice by day 5 of infection. As shown in Fig 5A, the groups receiving unadjuvanted QIV and QIV with K-Ac7-Dsa(- or SDI) LNPs showed less than 10% body weight loss over 5 days post infection. The unvaccinated PBS group lost approximately 20% body weight by 4DPI. Interestingly, the mice vaccinated with S-Ac7-Dog (-), irrespective of combination with IMDQ-PEG-Chol, showed drastic weight loss over 5 days, almost comparable to the unvaccinated PBS group. This is especially important to note because the same S- Ac7-Dog LNPs incorporating SDI did not show such extensive weight loss in vaccinated/challenged animals, suggesting higher morbidity in virus infected lungs in case of empty S-Ac7-Dog LNP formulations.

**Figure 5:**
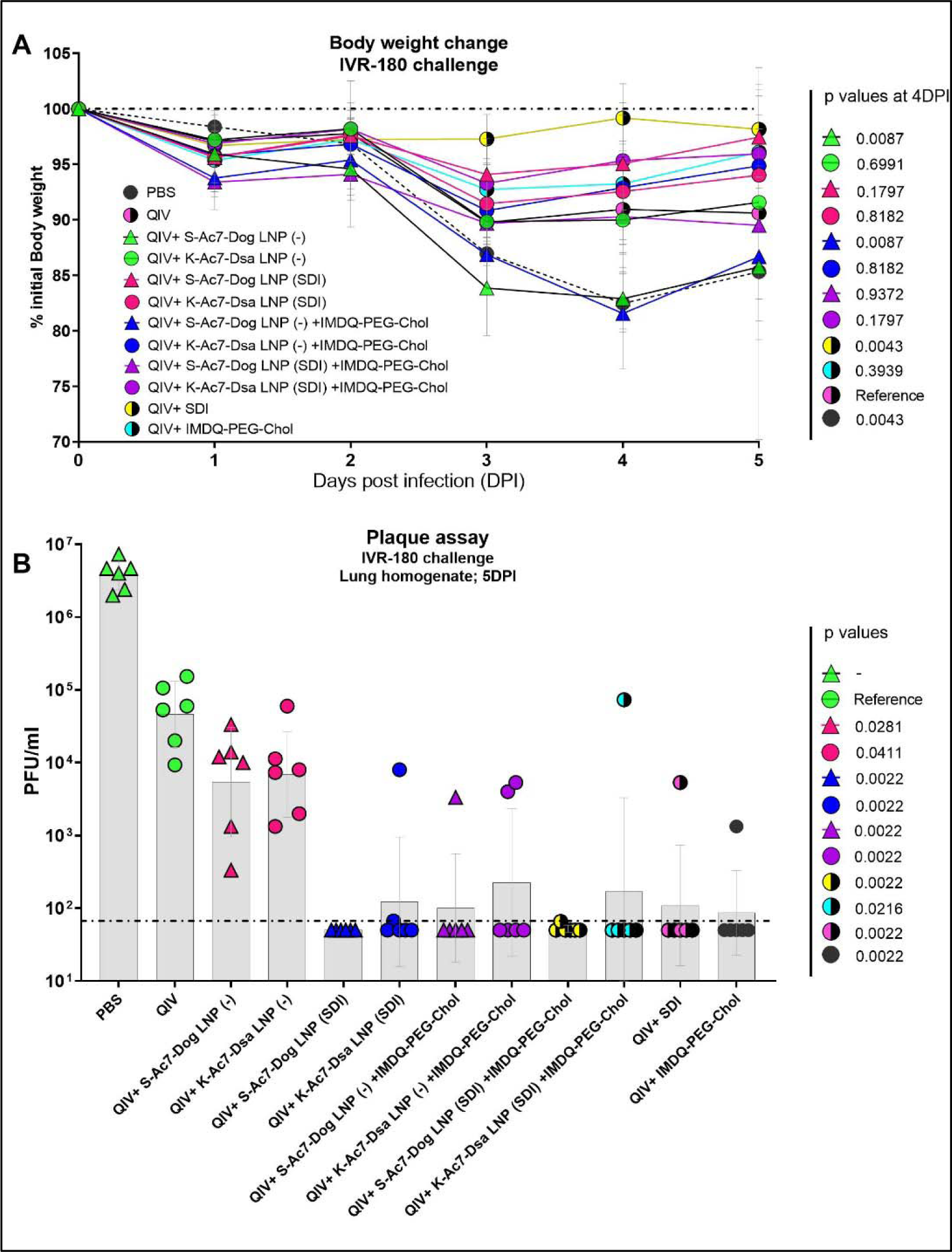
**QIV formulated in S-Ac7-Dog(SDI) and K-Ac7-Dsa(SDI) LNPs protects the vaccinated animals against a lethal dose of IVR-180, irrespective of combination with IMDQ- PEG-Chol**: All unvaccinated and QIV± IMDQ-PEG-Chol ± S-Ac7-Dog(SDI) or K-Ac7-Dsa(SDI) vaccinated animals were intranasally challenged with 100LD_50_ (18000 PFU per animal) of IVR-180 virus. **(A)** The body weight of each animal in all groups was recorded every day until the day of harvest and represented as percentage initial body weight for each group (n=6) (geometric mean ±geometric SD). **(B)** The viral lung titers were quantified at 5 days post infection by plaque assays on pre-seeded MDCK cells and are represented as PlaqueLformingLunit (PFU)/ml for n=6 animals per group (geometric mean ±geometric SD). Each data point represents one animal in the respective group. Statistical analysis was performed using two-sided Mann Whitney U test. The p values shown are calculated in reference to the unadjuvanted QIV group which received neither adjuvant nor LNP formulations.

Irrespective of the body weight loss differences attributed by LNPs, all groups which received adjuvanted QIV showed lower amount of replicating virus in their lungs 5 days post-infection (Fig 5B), compared with the unadjuvanted QIV group. QIV formulated in either of the two empty LNPs (S-Ac7-Dog(-) or K-Ac7-Dsa(-)) did not provide significant protection compared with unadjuvanted QIV group, contrary to the higher IgG1 induction. The groups that received QIV with either of S- Ac7-Dog(- or SDI) or K-Ac7-Dsa(- or SDI) LNPs, irrespective of IMDQ-PEG-Chol combination, resulted in significantly better control of lung virus replication, with very low detectable titers in their lungs and therefore, correlated with enhanced vaccine responses observed in mice. Interestingly, S-Ac7-Dog LNPs seemed to provide more protection compared to K-Ac7-Dsa LNPs in animals when compared between corresponding adjuvant groups.

### Incorporation of SDI as an adjuvant improves the cytokine profile in lungs of infected animals causing less morbidity as compared to empty S-Ac7-Dog LNP formulations

The body weight data over 5 days post infection suggested that the administration of empty S-Ac7- Dog LNP as vaccine ±adjuvant formulation was, although protective, did not provide protection from body weight loss compared to PBS-vaccinated control mice. The lungs of all infected animals were examined for their cytokine profiles 5 days post infection. As shown in Fig 6 and supplementary Fig 3, the cytokine profiles in the infected lungs of animals from vaccinated or unvaccinated/PBS groups were different. The PBS group showed high levels of inflammatory cytokines including IL-6, IL-18, IL-12 p70, IFN-γ, TNF-α as well as chemokine GMCSF which may suggest enhanced vascular permeability and increased infiltration of innate immune cells in response to infection in unvaccinated animals. As shown in Fig 6A, these cytokine levels were significantly lower in all vaccinated groups, implying a better control over inflammation and morbidity in response to the virus. Some of the cytokines were very low in PBS group, especially the signatures for T cell responses such as IL-4, IL-5 and IL-13, which is in line with the typical type 1 skewed host immune response to influenza infection. Interestingly, pro-inflammatory cytokines and chemokines, including IL-1β, GMCSF, and the type 2 cytokines IL-4, IL-5, IL-6, IL-13 were significantly elevated in unadjuvanted QIV and QIV± IMDQ-PEG-Chol formulated with empty LNPs. This was not the case for the corresponding groups with LNP(SDI), especially for those LNP groups with S- Ac7-Dog lipids. The PCA plot in Fig 6B clusters different vaccinated and unvaccinated groups based on their differences in production of inflammatory cytokines and chemokines in lungs post-infection. The unvaccinated PBS group clusters far away from all vaccinated animals corresponding to high inflammation and chemokine production in lungs post infection with lower interleukins. Besides, the QIV+ S-Ac7-Dog(-) vaccinated group clustered separately from the other vaccinated groups. Additionally, the group that received QIV+ S-Ac7-Dog(-) combined with IMDQ-PEG-Chol is clustered together with unadjuvanted QIV, suggesting that the use of empty S-Ac7-Dog is disadvantageous as it reduces the protective effect of IMDQ-PEG-Chol when used without any lipid formulation. The other vaccinated groups showing low inflammation in lungs clustered together.

**Figure 6:**
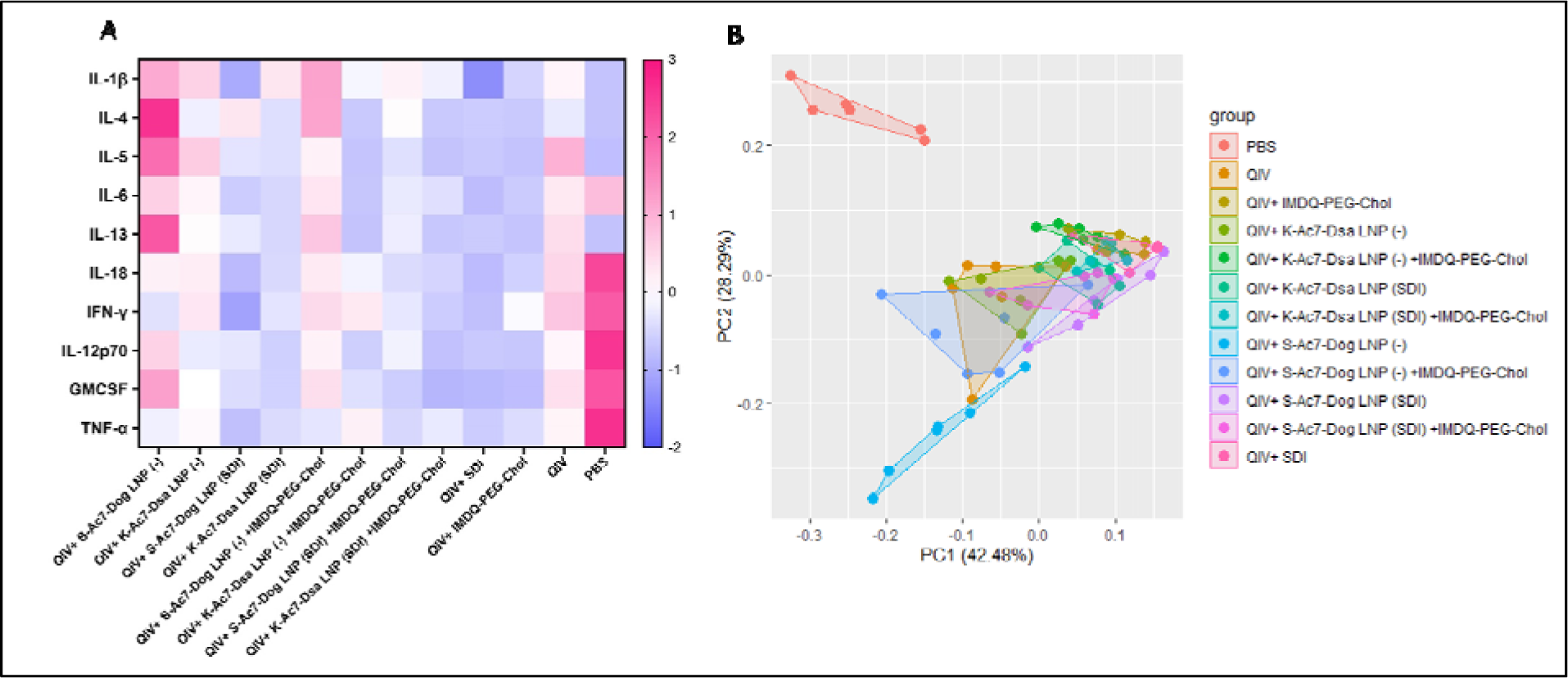
**Heatmap showing cytokine profile in lungs of vaccinated mice upon intranasal challenge with IVR-180**: All unvaccinated and vaccinated animals were intranasally challenged with 18000 PFU of IVR-180 virus. The lungs were collected at 5 days post infection and the cytokine levels were quantified by multiplex ELISA. Levels of IL-1β, IL-5, IL-13, TNF-α, IL-12 p70, IL-4, IL-6, IFN-γ, IL-18 and GMCSF, for n=6 animals per group are represented as median of z-scores (A) and PCA plots (B) for all animals in each group.

## Discussion

Quadrivalent inactivated influenza vaccines (QIV), consisting of two influenza A (IAV) and two influenza B (IBV) virus strain components and one of the licensed influenza vaccines, are modified and administered annually to provide immunity against circulating influenza viruses in human population. Several studies have been focussed on improving these split vaccines’ efficiency by combining them with adjuvants, including our recent study on RIG-I and TLR7/8 agonists as adjuvants for QIV. Then again, the stability of these vaccines ±adjuvants (such as SDI RNA; a RIG-I agonist) and their efficient *in vivo* delivery is still challenging. To overcome this, lipid nanoparticles (LNPs) have emerged as promising vehicles for *in vivo* delivery systems. Their ability to encapsulate, stably carry and efficiently deliver the molecules-of-interest has provided an effective platform in pharmaceutical as well as vaccine fields. One such example is highly efficient current mRNA vaccines for SARS-CoV-2 which use LNP-based formulations^24^. Nevertheless, these vaccines still need multiple booster doses to be sufficiently effective against emerging virus strains and therefore are limited in inducing an antigenically broader immune responses. Therefore, the LNPs doses and composition need to be tailored together with the vaccine and/or adjuvant combinations, in order to get the desired vaccination outcomes in terms of both humoral and cellular responses, besides providing protection against viral infections^22,25^.

In this study, we tested adjuvant formulations for QIV (2018-19 season; with A/ Singapore/GP1908/2015 IVR-180 (H1N1) as one of the two IAV components) with two different LNP formulations using either S-Ac7-Dog or K-Ac7-Dsa cationic lipids. Since the lipids are positively charged, they can stably interact with negatively charged biomolecules. The vaccine-LNP combinations were further adjuvanted with either one or both of RIG-I and TLR7/8 agonists (SDI- RNA and IMDQ-PEG-Chol, respectively), to explore the outcomes of the vaccine ±adjuvant ±LNP formulations in context of antibody responses, antibody class switching, T cell responses, protection against a lethal dose of IVR-180 virus and inflammatory responses in the infected lungs.

The unadjuvanted QIV vaccine induces modest serum IgG1 and IgG2a titres as well as low hemagglutinin inhibition titres 3 weeks after a single dose of intramuscular vaccination in mice. This is consistent with our previous findings using the same mouse vaccination model^17^. The low antibody titres for unadjuvanted QIV correlated well with low T cell responses (IL-4 and IFN-γ ELIspots) as well as inadequate protection against virus infection, implied by high levels of replicating virus in lungs upon intranasal IVR-180 challenge. These animals also showed most body weight loss and high levels of inflammatory cytokines in their lungs post infection, suggesting the recruitment of immune cells to aid in control/clearance of the virus. QIV formulated with either of two LNPs and/or further combined with either SDI or IMDQ-PEG-Chol as adjuvant, show a boosted total IgG response and a better control over virus replication by day 5 post infection. However, the administration of individual or combination adjuvants directed the B cell class switch as well as T cell responses in vaccinated mice. Upon a single vaccination, SDI induced a balanced IgG1/IgG2a response, IMDQ-PEG-Chol directed more towards an IgG2a response and the combination of the two skewed completely towards IgG2a, when formulated into S or K LNPs, suggesting very strong class switching events driven by this combination of adjuvants and consistent with our previous findings. Interestingly, the combination of the two adjuvants SDI and IMDQ-PEG-Chol induces a balanced type-I and -II T cell responses as suggested by a balanced IL-4 and IFN-γ release upon antigenic restimulation. Therefore, the combination of IMDQ-PEG-Chol and SDI in LNP formulations results in a balanced Th1/Th2 which results in complete type 1 skewing when antibody responses are considered.

Interestingly, S-Ac7-Dog LNPs were found to be more immunogenic than K-Ac7-Dsa LNPs in inducing both humoral and cellular responses in corresponding groups. Yet, vaccination with empty S-Ac7-Dog LNPs resulted in lack of protection from morbidity in challenged mice. Remarkably, S- Ac7-Dog(-) + IMDQ-PEG-Chol vaccination also resulted in lack of protection from morbidity after infection, with body weight loss comparable to unvaccinated/PBS control animals. This lack of protection from morbidity was accompanied by enhanced inflammatory cytokine responses including interleukins IL-4, IL-5, IL-6 and IL-13, interferon gamma (IFN-γ) as well as chemokine GMCSF. In contrast, when S-Ac7-Dog LNPs were combined with SDI, inflammation is relatively reduced in infected lungs as suggested by the chemokines/cytokines levels. The differences might plausibly be attributed by the lower stability of S-Ac7-Dog than K-Ac7-Dsa lipids in respective LNPs, which might be stabilized by addition of an opposite charged SDI molecules. However, we need additional experiments to confirm this.

Overall, we compared two different lipid compositions in LNP formulations, empty or loaded with individual or combination adjuvants. Different combinations affected both B and T cell responses along with vaccine/adjuvant/LNP-dependent inflammation in single vaccinated mice upon virus infection. The negatively charged SDI might stably interact with cationic lipid moieties providing more stability to the entire structure and thereby reducing related inflammation in infected animals. The immunogenicity and protection data in mice combined with the cytokine/chemokine induction indicates that lipid composition of LNPs used in vaccines is important and can skew host immune responses to subsequent infection, and therefore is important for vaccine safety and efficacy.

## Methods and Reagents

A list of reagents used in the study is provided in supplementary table I.

### Cell lines

Madin-Darby canine kidney (MDCK) cell line was maintained in Dulbecco’s Modified Eagle Medium (DMEM) supplemented with 10% Fetal bovine serum (FBS) and 1X antibiotics (penicillin/ streptomycin).

### QIV vaccine

Quadrivalent Inactivated influenza vaccine (FLUCALVEX 2018/2019 season Lot 252681) was obtained from Seqirus. The vaccine consists of MDCK-grown two Influenza A and two B viruses- A/ Singapore/GP1908/2015 IVR-180 (H1N1) (A/Michigan/45/ 2015-like virus), A/North Carolina/04/2016 (H3N2) (A/ Singapore/INFIMH-16-0019/2016 -like virus), B/Iowa/06/2017 (B/Colorado/06/2017-like virus) and B/Singapore/INFTT-16- 0610/2016 (B/Phuket/3073/2013-like virus).

### SDI-RNA

The SDI-RNA (or SDI) was *in vitro* transcribed as described in our recent study.

### Lipid nanoparticle (LNP) fabrication

1μg SDI or SDI equivalent was encapsulated in LNPs by rapid mixing under vigorous stirring of an acetate buffer (5 mM, pH 4.5) containing SDI with an ethanolic solution containing the ionizable lipid S-Ac7-Dog or K-Ac7-Dsa (to obtain S-Ac7-Dog and K-Ac7-Dsa LNPs respectively), cholesterol, 1,2-dioleoyl-n-glycero-3-phosphoethanolamine (DOPE) and a poly(ethylene glycol)- lipid (DSG-PEG; PEG had an MW of 2 kDa) at a 50:38.5:10:1.5 ratio. After mixing, LNP were dialyzed against 1X PBS to get rid of ethanol and the pH was adjusted to 7.4. An N/P (ionizable Nitrogen atoms of the ionizable lipid to anionic Phosphor atoms of SDI) molar ratio of 5:1 was targeted.

### LNP characterization

The diameter and polydispersity index (PDI) of LNPs was measured with Dynamic light scattering (DLS) at physiological pH. Each sample was measured in triplicates and a cumulative average of z average and PDI was calculated. For ELS, each sample was appropriately diluted in HEPES buffer and measurements were taken in triplicates. The Zeta potential was calculated for all samples based on the Smoluchowski equation.

### Vaccine-adjuvant preparation and administration

For each animal, 1.5mg HA equivalent of QIV was mixed with empty or SDI-encapsulating S-Ac7- Dog or K-Ac7-Dsa LNPs, with or without 100μg IMDQ-PEG-Chol (equivalent to 10μg core IMDQ), and vortexed for 30 seconds (s). Adjuvant doses were chosen based on our previously published work with these adjuvants^17,30^. Unadjuvanted or adjuvanted QIV, with or without LNP formulations, was administered intramuscularly in a total of 50μl per mouse, in the right hind leg. The control group were administered with equal volume of PBS instead of vaccine or vaccine ± adjuvant ± LNP mixture. All animals received only one dose of vaccine without any further boosters.

### IVR-180 virus

A/Singapore/GP1908/2015 IVR-180 (H1N1) was grown in 8-days old embryonated chicken eggs and was titrated by plaque assay on pre-seeded MDCK cells. The half-lethal dose of virus needed to kill mice (LD_50_) was calculated based on intranasal infections in 6-8 weeks old naïve female BALB/c mice using 10-fold serial dilutions of the virus.

### Mouse model

The study was performed on 6–8-week-old female BALB/c mice strains obtained from Charles River Laboratories, MA. The mice were housed with food and water ad libitum in a pathogen-free facility at Icahn School of Medicine at Mount Sinai, New York. All mice were vaccinated intramuscularly (50μl; right hind leg; per mouse) and infected intranasally (in 50μl total volume per mouse) under ketamine/xylazine anesthesia. All procedures were performed according to the protocols approved by the Icahn School of Medicine at Mount Sinai Institutional Animal Care and Use Committee (IACUC -PROTO202100007).

### Serum collection for serology

Mice blood was collected by submandibular bleed 3 weeks post vaccination from all animals. The blood was allowed to clot at 4°C for overnight. The serum was collected after a brief centrifugation and was heat inactivated at 56°C for 30 min. The samples were stored at -20°C until further use.

### Enzyme-linked immunosorbent assay

Enzyme-linked immunosorbent assay (ELISA) was performed to quantify the vaccine-specific IgG titers in mice sera. Briefly, ELISA plates were coated with recombinant trimeric HA (derived from the A/Michigan/45/2015 H1N1 virus, which is closely related to IVR-180; as previously described^17^, equivalent to 2µg H1N1-HA/ml, in bicarbonate buffer and left overnight at 4°C. Plates were washed three times with 1X PBS and incubated in 100μl blocking buffer per well (5% fat-free milk in PBST (1X PBS + 0.1% Tween20)) for 1 hour (h) at room temperature (RT). In the meantime, the serum samples were serially diluted 3-fold, starting with 1:100 dilution, in the blocking buffer and 50μl of each sample dilution was incubated on HA-coated ELISA plates for overnight at 4°C. The following day, the plates were washed three times with PBST and incubated with 100μl of diluted horse radish peroxidase (HRP)-conjugated anti-mouse secondary total IgG (1:5000) or IgG1 (1:2000) or IgG2a (1:2000) antibodies, for 1h at RT. Finally, the plates were washed three times in PBST and incubated with 100µl of tetramethylbenzidine (TMB) substrate at RT until the blue color appeared. The reaction was terminated with 50µl of 1M sulfuric acid (H_2_SO_4_), and the absorbance was measured at 450nm and 650nm wavelengths using BIOTEK ELISA plate reader.

### Hemagglutination inhibition assay (HAI)

Mice sera collected 3 weeks post-vaccination were treated with 4 volumes of receptor destroying enzyme (RDE) at 37°C overnight, followed by treatment with 5 volumes of 1.5% sodium citrate at 56°C, 30 min. Thus obtained 1:20-diluted serum samples were further serially diluted in a transparent V-bottom 96-well plate and incubated with 4HA units per well of IVR-180 virus for 30 min at RT, followed by addition of 0.5% chicken red blood cells for 40 min at 4°C. The results were recorded as HAI titers.

### Enzyme-linked immunosorbent spot

Mice were vaccinated with different combinations of QIV ±adjuvant ±LNPs and the spleens were harvested 10 days post-vaccination from all vaccinated as well as unvaccinated animals. The spleens were collected in 5ml RPMI-1640 media supplemented with 2% FBS and 1X penicillin/streptomycin and kept on ice. A single cell suspension of splenocytes was obtained by homogenizing the spleens against a 70μm strainer. Interferon gamma (IFN-γ) or Interleukin-4 (IL-4) enzyme-linked immunosorbent spot (ELIspot) assays were performed using 10^5^ splenocytes per well in a 96-well Polyvinylidene difluoride (PVDF) ELIspot plates, precoated with IFN-γ or IL-4 capture antibodies, respectively, according to manufacturer’s protocol. Splenocytes were unstimulated or restimulated either with hemagglutinin (HA-H1N1) overlapping 15-mer peptides or whole live IVR-180 (H1N1) virus and incubated overnight in 37°C incubator. The wells were washed thrice with water to get rid of cells and incubated with 100μl of biotinylated polyclonal detection antibody against IFN-γ or IL-4 for 1.5h at RT, followed by an incubation with streptavidin-HRP conjugated antibody for 1h at RT. The plates were finally incubated with 100μl of the substrate for 1h in dark, followed by thorough washing under tap water multiple times. The plates were air dried in dark and the number of spots in each well were manually counted using a dissection microscope. The results were represented as number of IFN-γ or IL-4 producing splenocytes per million splenocytes.

### Virus challenge

100LD_50_ of IVR-180 (18000 plaque forming units (PFU) per animal), was used for intranasal infection in a final volume of 50µL per mouse. The virus challenge was performed under mild anesthesia with ketamin/xylazine (intraperitoneal) as recommended by AICUC. The unvaccinated but challenged group was used as a control in the experiment. Body weights were recorded every day post-infection until lung harvest. The lungs were collected at 5 days post-infection (DPI) in 500μl 1X PBS and homogenized using a tissue homogenizer. The lysate thus obtained was stored at -80°C for viral titrations.

### Plaque assay

Virus titrations were performed by plaque assays to quantify the replicating virus titers in the lungs of vaccinated versus unvaccinated mice. The lung homogenate (or lysate) was 10-fold serially diluted in 1X PBS and incubated on pre-seeded and pre-washed monolayers of MDCK cells for 1h in an incubator, at 37°C, 5% CO_2_ with gentle shaking every 5 min. The diluted samples were then removed, and the monolayers were again briefly washed with 1ml 1X PBS. Lastly, 1ml of the overlay mixture (2% oxoid agar and 2X minimal essential medium (MEM) supplemented with 1% diethyl-aminoethyl (DEAE)-dextran and 1 mg/ml tosylamide-2- phenylethyl chloromethyl ketone (TPCK)-treated trypsin) was added on top of the monolayers and incubated for 48 h in the incubator, at 37°C, 5% CO_2_. The plates were finally fixed in 4% formaldehyde. The overlay was removed, and the plaques were immune-stained with IVR-180-postchallenge polyclonal serum, diluted 1:1000 in blocking buffer. The plates were washed and incubated with 1:5000 dilution of horse radish peroxidase (HRP)-conjugated anti-mouse secondary antibody for 1h at RT with gentle shaking. Followed by a brief washing in 1X PBST, the plaques were finally visualized with True Blue substrate and the number of plaques were counted and represented as PFU/ml.

### Multiplex ELISA

Multiplex ELISA was performed for simultaneous measurements of different cytokines in the lung homogenates from IVR-180-infected mice. The following cytokines were measured: Granulocyte macrophage colony-stimulating factor (GMCSF), Interleukin (IL)-1β, IL-4, IL-5, IL6, IL-12p70, IL- 13, IL-18 and Interferon gamma (IFN-γ). The assay was performed according to the manufacturer’s guidelines and the readings were recorded using Luminex 100/200 system.

### Software

The schematic figures were created with BioRender.com. GraphPad Prism version 10 was used for data visualization, analysis, graph plotting and statistical analysis.

### Data availability statement

The original contributions in the study are included in the article/supplementary material. Further inquiries can be directed to the corresponding author at.

## Ethics statement

The animal study was reviewed and approved by the Institutional Animal Care and Use Committee (IACUC) at the Icahn School of Medicine at Mount Sinai, New York.

## Author contributions

Conceptualization and study design: SJ and MS.

Methodology: SJ (viruses, infections, mouse immunization), AL, YC and TY (LNP fabrication and characterization), SJ and GL (*in vitro* serological assays), SJ and GS (ELIspots), SJ (multiplex ELISA).

Reagents: SJ, AL, TY, YC, AG-S, BG and MS.

Investigation and data analysis: SJ and MS. First draft of the manuscript: SJ and MS. Manuscript review and editing: all authors. Funding acquisition: AG-S, BG, and MS.

All authors contributed to the article and approved the submitted version.

## Funding and Acknowledgement

This study was partly funded by CRIPT (Center for Research on Influenza Pathogenesis and Transmission), a NIH NIAID funded Center of Excellence for Influenza Research and Response (CEIRR, contract number 75N93021C00014), by the NIAID funded SEM-CIVIC consortium to improve influenza vaccines (contract number 75N93019C00051) and by NIAID grant P01AI097092 to AG-S. Influenza research in the M.S. lab is further supported by NIH/NIAID R21AI151229, R44AI176894 and R21AI176069. BG acknowledges funding from the European Research Council (ERC) under the European Union’s Horizon 2020 research and innovation program (grant N 817938).

## Supplementary Table

## Supplementary Figures

**Supplementary figure 1:**
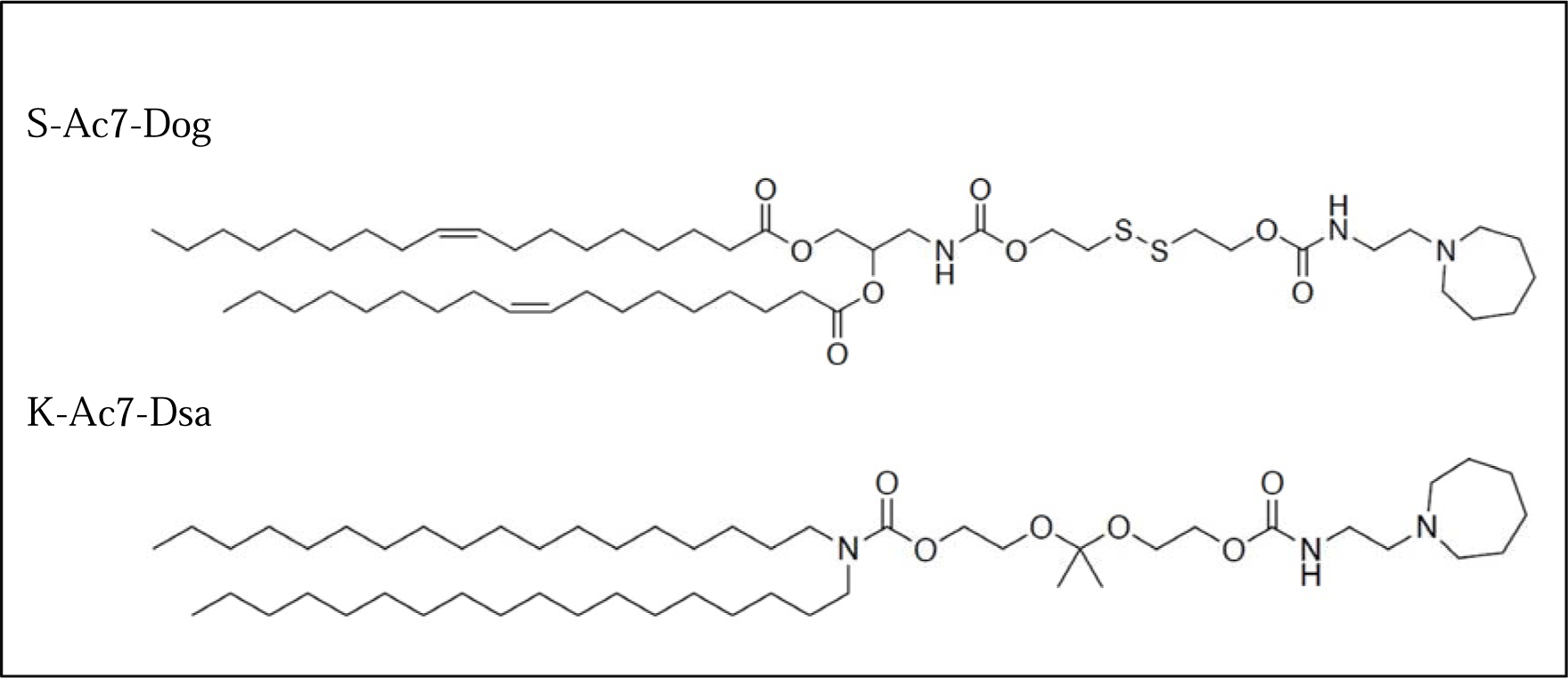
Chemical structure of in-house synthesized ionizable lipids- S-Ac7-Dog and K-Ac7-Dsa lipids, comprising a disulfide bond that can be cleaved by reduction and a ketal bond that can be cleaved by acidic pH, respectively.

**Supplementary figure 2:**
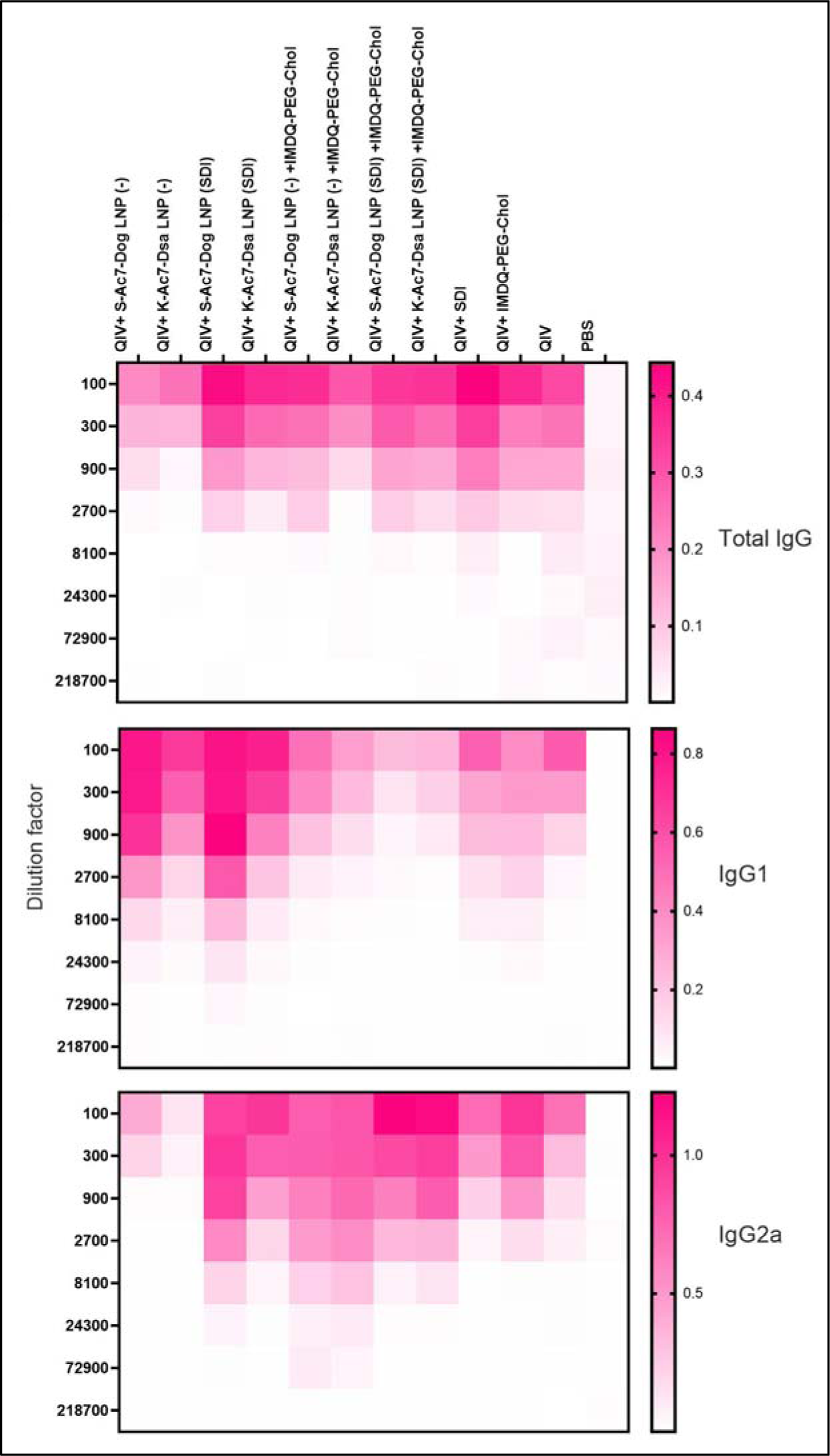
Heatmap showing mean OD450 ELISA values (n=5 per group) plotted against serum dilutions for total IgG, IgG1 and IgG2a. 6-8-week female BALB/c mice were vaccinated with QIV with and without IMDQ-PEG-Chol and formulated into empty or SDI- encapsulating S or K LNPs. Serum was collected 3 weeks post-vaccination by submandibular bleed. Total IgG, IgG1 and IgG2a titers were quantified by ELISA with 3-fold serum dilutions starting with 1:100, for H1 HA specific antibodies.

**Supplementary figure 3:**
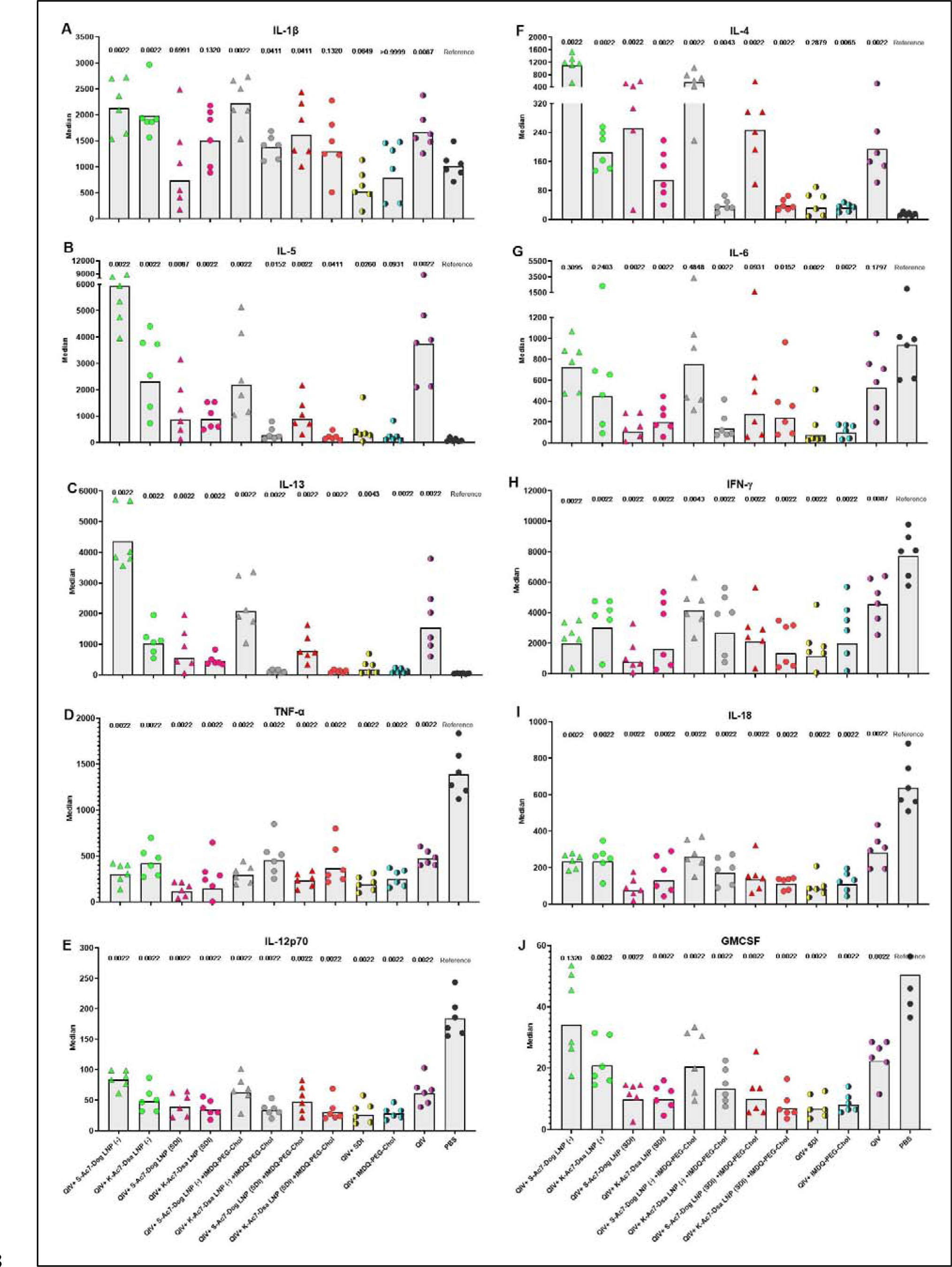
All lungs from unvaccinated and vaccinated animals were collected at 5 days post infection and the cytokine levels were quantified by multiplex ELISA. Levels of **(A)** IL- 1β, **(B)** IL-5, **(C)** IL-13, **(D)** TNF-α, **(E)** IL-12 p70, **(F)** IL-4, **(G)** IL-6, **(H)** IFN-γ, **(I)** IL-18 and **(J)** GMCSF, for n=6 animals per group are represented as geometric mean ±geometric SD, where each data point corresponds to individual mouse. Statistical analysis was performed using two-sided Mann Whitney U test. The p values shown are calculated in reference to the virus-challenged unvaccinated group (denoted as PBS).

**Supplementary table I:** List of reagents and kits used in the study.

| Reagent | Brand | Catalogue number |
| --- | --- | --- |
| DMEM | Corning | 10013-CV |
| RPMI 1640 | Gibco | 22400089 |
| Penicillin/streptomycin | Corning | 30002-CI |
| Goat HRP-conjugated secondary Anti-mouse IgG antibody | Abcam | Ab6823 |
| Anti-mouse IgG1-HRP secondary antibody | Invitrogen | PA174421 |
| Anti-mouse IgG2a-HRP secondary antibody | Invitrogen | A10685 |
| TMB substrate | BD OptEIA | 555214 |
| KPL true blue substrate | Sera care | 5510-0050 |
| 70µm strainer | BD | 352340 |
| Peptivator H1-HA peptide | MiltenyiBiotec | 130-099-803 |
| Oxoid agar | Thermofisher | LP0028B |
| EMEM | BioWhittaker | 12684F |
| TPCK | Sigma | T4376 |
| 10% methanol-free formaldehyde | Polysciences | 040181 |
| 2M Sulfuric acid | Thermo fisher | S25898 |
| Mouse IFN-γ ELISPOT kit | RnD systems | EL485 |
| Mouse IL-4 ELISPOT kit | RnD systems | EL404 |
| ELISA NUNC-maxisorp plates | Invitrogen | 44240421 |
| Th1/Th2 Cytokine 11-Plex Mouse ProcartaPlex™ Panel | Invitrogen | EPX110-20820-901 |

## References

1. Cox RJ, Brokstad KA, Ogra P. Influenza virus: immunity and vaccination strategies. Comparison of the immune response to inactivated and live, attenuated influenza vaccines. Scand J Immunol. 2004;59(1):1–15. doi:10.1111/j.0300-9475.2004.01382.x

2. Hannoun C. The evolving history of influenza viruses and influenza vaccines. Expert Rev Vaccines. 2013;12(9):1085–1094. doi:10.1586/14760584.2013.824709

3. Kim H, Webster RG, Webby RJ. Influenza Virus: Dealing with a Drifting and Shifting Pathogen. Viral Immunol. 2018;31(2):174–183. doi:10.1089/vim.2017.0141

4. Boni MF. Vaccination and antigenic drift in influenza. Vaccine. 2008;26 Suppl 3(Suppl 3):C8-14. doi:10.1016/j.vaccine.2008.04.011

5. De Jong JC, Rimmelzwaan GF, Fouchier RA, Osterhaus AD. Influenza virus: a master of metamorphosis. J Infect. 2000;40(3):218–228. doi:10.1053/jinf.2000.0652

6. Mooij P, Mortier D, Aartse A, et al. Vaccine-induced neutralizing antibody responses to seasonal influenza virus H1N1 strains are not enhanced during subsequent pandemic H1N1 infection. Front Immunol. 2023;14:1256094. doi:10.3389/fimmu.2023.1256094

7. Petrie JG, Ohmit SE, Truscon R, et al. Modest Waning of Influenza Vaccine Efficacy and Antibody Titers During the 2007-2008 Influenza Season. J Infect Dis. 2016;214(8):1142–1149. doi:10.1093/infdis/jiw105

8. Hsu JP, Zhao X, Chen MIC, et al. Rate of decline of antibody titers to pandemic influenza A (H1N1-2009) by hemagglutination inhibition and virus microneutralization assays in a cohort of seroconverting adults in Singapore. BMC Infect Dis. 2014;14(1):414. doi:10.1186/1471-2334-14-414

9. Palese P, García-Sastre A. Influenza vaccines: present and future. J Clin Invest. 2002;110(1):9-13. doi:10.1172/JCI15999

10. Harding AT, Heaton NS. Efforts to Improve the Seasonal Influenza Vaccine. Vaccines (Basel). 2018;6(2):19. doi:10.3390/vaccines6020019

11. Coutelier JP, van der Logt JT, Heessen FW, Warnier G, Van Snick J. IgG2a restriction of murine antibodies elicited by viral infections. J Exp Med. 1987;165(1):64–69. doi:10.1084/jem.165.1.64

12. Coutelier JP, van der Logt JT, Heessen FW, Vink A, van Snick J. Virally induced modulation of murine IgG antibody subclasses. J Exp Med. 1988;168(6):2373–2378. doi:10.1084/jem.168.6.2373

13. Hovden AO, Cox RJ, Haaheim LR. Whole influenza virus vaccine is more immunogenic than split influenza virus vaccine and induces primarily an IgG2a response in BALB/c mice. Scand J Immunol. 2005;62(1):36–44. doi:10.1111/j.1365-3083.2005.01633.x

14. Barackman JD, Ott G, O’Hagan DT. Intranasal immunization of mice with influenza vaccine in combination with the adjuvant LT-R72 induces potent mucosal and serum immunity which is stronger than that with traditional intramuscular immunization. Infect Immun. 1999;67(8):4276–4279. doi:10.1128/IAI.67.8.4276-4279.1999

15. Moran TM, Park H, Fernandez-Sesma A, Schulman JL. Th2 responses to inactivated influenza virus can Be converted to Th1 responses and facilitate recovery from heterosubtypic virus infection. J Infect Dis. 1999;180(3):579–585. doi:10.1086/314952

16. Hovden AO, Cox RJ, Madhun A, Haaheim LR. Two doses of parenterally administered split influenza virus vaccine elicited high serum IgG concentrations which effectively limited viral shedding upon challenge in mice. Scand J Immunol. 2005;62(4):342–352. doi:10.1111/j.1365-3083.2005.01666.x

17. Jangra S, Laghlali G, Choi A, et al. RIG-I and TLR-7/8 agonists as combination adjuvant shapes unique antibody and cellular vaccine responses to seasonal influenza vaccine. Front Immunol. 2022;13:974016. doi:10.3389/fimmu.2022.974016

18. Visciano ML, Tagliamonte M, Tornesello ML, Buonaguro FM, Buonaguro L. Effects of adjuvants on IgG subclasses elicited by virus-like particles. J Transl Med. 2012;10:4. doi:10.1186/1479-5876-10-4

19. Huber VC, McKeon RM, Brackin MN, et al. Distinct contributions of vaccine-induced immunoglobulin G1 (IgG1) and IgG2a antibodies to protective immunity against influenza. Clin Vaccine Immunol. 2006;13(9):981–990. doi:10.1128/CVI.00156-06

20. Pulendran B, S Arunachalam P, O’Hagan DT. Emerging concepts in the science of vaccine adjuvants. Nat Rev Drug Discov. 2021;20(6):454–475. doi:10.1038/s41573-021-00163-y

21. Snapper CM, Paul WE. Interferon-gamma and B cell stimulatory factor-1 reciprocally regulate Ig isotype production. Science. 1987;236(4804):944-947. doi:10.1126/science.3107127

22. Lee Y, Jeong M, Park J, Jung H, Lee H. Immunogenicity of lipid nanoparticles and its impact on the efficacy of mRNA vaccines and therapeutics. Exp Mol Med. 2023;55(10):2085–2096. doi:10.1038/s12276-023-01086-x

23. Tenchov R, Bird R, Curtze AE, Zhou Q. Lipid Nanoparticles─From Liposomes to mRNA Vaccine Delivery, a Landscape of Research Diversity and Advancement. ACS Nano. 2021;15(11):16982–17015. doi:10.1021/acsnano.1c04996

24. Schoenmaker L, Witzigmann D, Kulkarni JA, et al. mRNA-lipid nanoparticle COVID-19 vaccines: Structure and stability. Int J Pharm. 2021;601:120586. doi:10.1016/j.ijpharm.2021.120586

25. Zhong Z, Chen Y, Deswarte K, et al. Lipid Nanoparticle Delivery Alters the Adjuvanticity of the TLR9 Agonist CpG by Innate Immune Activation in Lymphoid Tissue. Adv Healthc Mater. Published online September 29, 2023:e2301687. doi:10.1002/adhm.202301687

26. Zhang C, Ma Y, Zhang J, et al. Modification of Lipid-Based Nanoparticles: An Efficient Delivery System for Nucleic Acid-Based Immunotherapy. Molecules. 2022;27(6):1943. doi:10.3390/molecules27061943

27. Lamoot A, Jangra S, Laghlali G, et al. Lipid Nanoparticle Encapsulation Empowers Poly(I:C) to Activate Cytoplasmic RLRs and Thereby Increases Its Adjuvanticity. Small. Published online October 22, 2023:2306892. doi:10.1002/smll.202306892

28. Martínez-Gil L, Goff PH, Hai R, García-Sastre A, Shaw ML, Palese P. A Sendai virus-derived RNA agonist of RIG-I as a virus vaccine adjuvant. J Virol. 2013;87(3):1290–1300. doi:10.1128/JVI.02338-12

29. Patel JR, Jain A, Chou Y ying, Baum A, Ha T, García-Sastre A. ATPase-driven oligomerization of RIG-I on RNA allows optimal activation of type-I interferon. EMBO Rep. 2013;14(9):780–787. doi:10.1038/embor.2013.102

30. Jangra S, De Vrieze J, Choi A, et al. Sterilizing Immunity against SARS-CoV-2 Infection in Mice by a Single-Shot and Modified Imidazoquinoline TLR7/8 Agonist-Adjuvanted Recombinant Spike Protein Vaccine. bioRxiv. Published online October 23, 2020:2020.10.23.344085. doi:10.1101/2020.10.23.344085

